# Stiffening Symphony of Aging: How Senescent Osteocytes Lose Their Elastic Rhythm

**DOI:** 10.1101/2024.09.28.615588

**Authors:** Maryam Tilton, Megan Weivoda, Maria Astudillo Potes, Anne Gingery, Alan Y. Liu, Tamara Tchkonia, Lichun Lu, James L. Kirkland

## Abstract

Senescent osteocytes are key contributors to age-related bone loss and fragility; however, the impact of mechanobiological changes in these cells remains poorly understood. This study provides a novel analysis of these changes in primary osteocytes following irradiation-induced senescence. By integrating sub-cellular mechanical measurements with gene expression analyses, we identified significant, time-dependent alterations in the mechanical properties of these cells. Increases in SA-β-Gal activity and p16Ink4a expression levels confirmed the senescence status post-irradiation. Key findings include a time-dependent increase in cytoskeletal Young’s modulus and altered viscoelastic properties of the plasma membrane, affecting the contractility of primary osteocytes. Additionally, we observed a significant increase in Sclerostin (Sost) expression 21 days post-irradiation. These mechanobiological changes may impair osteocyte mechanosensation and mechanotransduction, contributing to bone fragility. This is the first study to time-map senescence-associated mechanical changes in the osteocyte cytoskeleton. Our findings highlight the potential of biophysical markers as indicators of cellular senescence, providing more specificity than traditional, variable biomolecular markers. We believe these results support biomechanical stimulation as a potential therapeutic strategy to rejuvenate aging osteocytes and enhance bone health.

## Introduction, Results, and Discussion

Cellular senescence, a hallmark of aging, is linked to a wide range of disorders and diseases, including neurodegenerative conditions, cancer, and degenerative musculoskeletal diseases such as bone loss.^1–5^ In the United States, over 43 million individuals aged 50 and older have low bone mass, placing them at high risk for osteoporosis and fragility fractures. ^6,7^ The prevalence of bone loss is notably higher in women compared to men. ^6,7^ With the global aging population, the economic burden of age-related fragility fractures is projected to exceed $25 billion by 2025.^8^ This underscores the critical need to understand the mechanobiological mechanisms underlying age-related bone loss. Senescent cells (SnCs), including senescent osteocytes, accumulate with age in various tissues, including bone. ^9–12^ These cells exhibit a senescence-associated secretory phenotype (SASP), characterized by the secretion of pro-inflammatory cytokines, chemokines, growth factors, proteases, and other factors, which can damage local and distant tissues and promote chronic inflammation. ^9,11–14^

Osteocytes, once thought to be mere “passive placeholders” in mineralized bone, have emerged as crucial multifunctional cells.^15–17^ These “superstars”^15^ play several essential roles: they are master regulators of bone homeostasis, endocrine cells that regulate phosphate metabolism in organs such as the kidney and parathyroid, and most importantly, they act as strain gauges, regulating bone mechanosensation and mechanotransduction. ^15–17^ The cytoskeleton of osteocytes, composed of actin microfilaments, microtubules, and intermediate filaments, is crucial for their mechanosensing capabilities.^15,16,18^ Mechanical stimulation induces cytoskeletal rearrangements and stress fiber formation, which are essential for the proper function of osteocytes. ^15,16,18^ However, with aging, the structure and mechanics of the cytoskeleton are altered, impairing osteocyte function and diminishing their mechanosensory properties.^18^ This dysfunction is exacerbated by the accumulation of SnCs and their SASP, both locally and systemically.^19^ These cells express markers of senescence, such as the cyclin-dependent kinase inhibitors p16^Ink4a^ (encoded by Cdkn2a) and p21^Cip1^ (encoded by Cdkn1a), which are part of the stress response program that drives cellular aging.^20^

Current research primarily relies on SA-β-Gal activity, expression of senescence and senescence-associated genomic markers, chromosomal changes, as well as combination of SA-β-Gal and flow cytometry to identify senescence status.^1,4,21–23^ However, these biomarkers are highly variable across different cell types and tissues.^1,21^ Additionally, there is evidence of sex-dependent variations in senescence-associated factors.^1,3,4,21,22^ Therefore, it is crucial to explore alternative biophysical markers that provide unique and reliable fingerprints of aging. Understanding changes in cytoskeletal mechanics could lay the foundation for future research aimed at reversing or rejuvenating aging cells through biomechanical stimulation. Moreover, establishing such biophysical markers could complement the growing set of genetic markers in this research space,^21^ offering more consistent and robust tools for identifying senescence and reducing reliance on the highly variable biomolecular markers currently in use.

Here, we present studies of novel biophysical markers of senescent primary osteocytes, focusing on changes in the mechanical properties of the plasma membrane and cytoskeleton following irradiation-induced senescence. By characterizing these biophysical alterations, we aim to establish reliable markers that can enhance understanding of cellular senescence and support the development of therapeutic strategies to counteract the effects of aging on bone health and potentially other tissues and organs.

Primary osteocytes were isolated from the vertebrae of 10 young (4-months) female C57BL/6 WT mice following our previously established protocols (Figure 1A).^4,20,22^ We used low-dose irradiation (10 Gy) as a pro-senescence stressor to induce senescence in primary osteocyte cultures *in vitro*. The isolated primary cells were cultured in α-MEM growth medium supplemented with 5% fetal bovine serum and 5% calf serum for 7 days prior to passaging into multiple 6-well plates for the experiments. All culture plates were coated with rat tail type I collagen (0.15 mg/mL). Maintaining three wells per study group for every experiment, we investigated changes in single-cell mechanics at different time points post-irradiation (D7, D14, and D21) and compared them to a non-senescent control group (CTRL). We used optical fiber-based interferometry nanoindentation (Pavone, Optics11Life) with an indenter tip size of R=3 µm and stiffness of 0.019 N/m on live cells, enabling time-lapsed single-cell analyses of senescence-associated changes in cytoskeletal modulus and membrane viscoelasticity. We included a minimum of 30 cells per well in our single-cell nanoindentation experiments, ensuring a minimum of 90 cells per study group per time point were tested, distributed across multiple wells. The senescence status of the cultures was determined by SA-β-Gal activity (Figure 1B). Real-Time quantitative Polymerase Chain Reaction (RT-qPCR) was performed to measure changes in the expression of senescence markers (*e*.*g*., p16^Ink4a^ and p21^Cip1^) (Figure 1C). Additionally, key genomic markers associated with osteocytes, Wnt/β-catenin signaling pathway, and mineralization, such as Sclerostin (Sost), Matrix Extracellular Phosphoglycoprotein (Mepe), and Matrix Metalloproteinases (Mmps), were measured using RT-qPCR (Figure 1C-D).

**Figure 1:**
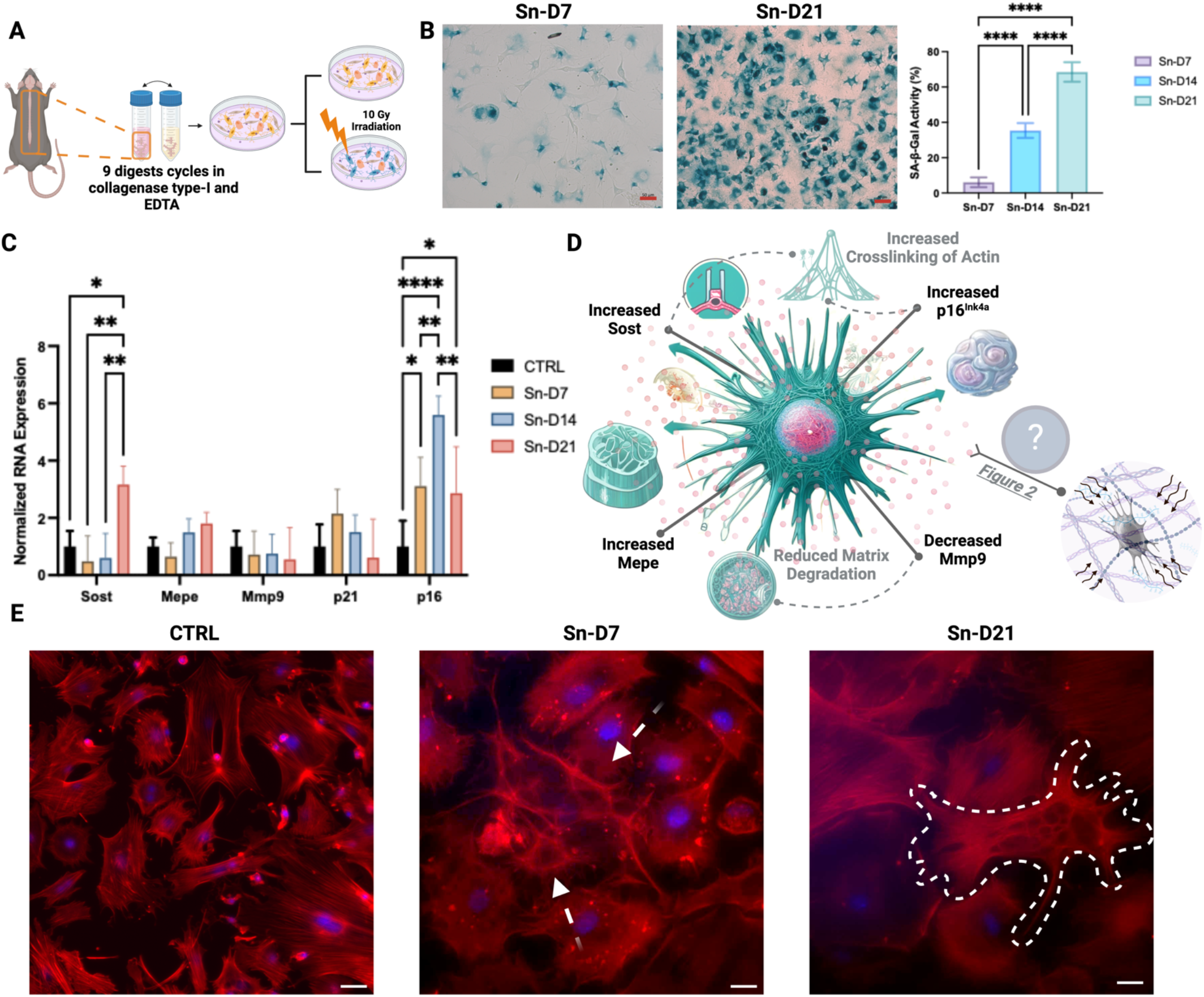
Overview of the study and gene expression changes in senescent primary osteocytes. (A) Schematic representation of primary osteocyte isolation, culture, and induction of senescence by irradiation. (B) Bright field microscopy of irradiated primary cell culture showing increased in SA-β-Gal activity with peaking to over 70% by day 21 [scale bar = 50 µm]. (C-D) Normalized RNA expression levels of key osteocyte and senescence-associated genes. Increases in Sost and Mepe, along with a decrease in Mmp9, were observed in senescent osteocytes compared to non-senescent control cells. Elevated p16 expression, along with increased SA-β-Gal activity (B), confirmed the senescence status. As illustrated in representative immunofluorescence staining images (F-actin/DAPI) [scale bar = 25 µm], (E) changes in cell size, dendritic network (arrows in Sn-D7), and morphology (dashed boundary in Sn-D21) were observed. We have provided additional IF images in the Supplementary Methods. These micrographs, from different regions of the CTRL and Sn-D21 cultures, allow for better observation of the morphological changes due to senescence condition across different regions. Data are presented as means ± SD, with statistical significance denoted by *p < 0.05, **p < 0.01, ***p < 0.001, and ****p < 0.0001.

As seen in Figure 2, using Hertzian contact mechanics model with our load-indentation data, ^24,25^ our findings indicated a significant increase in Young’s modulus of cells on D7, D14, and D21 post-irradiation compared to the CTRL group. Although the changes in modulus from D7 to D14 and D14 to D21 were not statistically significant, an increasing trend was observed, with a statistically significant increase at D21 compared to D7 (Figure 2B). Dissecting the unloading portion of indentation curves (Figure 2C) revealed a smoother recovery phase with a low adhesion force (range: 0.002 – 0.005 µN) in the CTRL groups, indicating that healthy primary cells exhibit predominantly elastic behavior. In contrast, senescent cells exhibited higher minimum adhesion force (range: 0.006 – 0.01 µN) with a noticeable secondary peak or ‘bump’ before returning to the baseline load. Analyzing the unloading portion of load-time graphs (Figure 2D), healthy cells showed smooth and rapid recovery to the baseline load, likely due to better stress relaxation capabilities. However, in senescent cells, the fluctuations (i.e., secondary peak) after the minimum adhesion force and the extended recovery time suggest heterogeneous viscoelastic properties and delayed mechanical recovery, indicating altered cytoskeletal architecture and function, a hallmark of senescence. Specifically, the wider interval between the minimum adhesion force and the return to the baseline load in senescent cells demonstrates the prolonged interaction between the indenter and the cells due to altered viscoelastic properties. These observations mark distinct biophysical fingerprints of cellular senescence. A previous study^26^ on senescent mesenchymal stem cells showed that irradiation-induced senescence led to cytoskeletal re-organization through formation of stress fibers, increased crosslinking of actin filaments, and a denser actin network. The cytoskeletal stiffening process increases the cells’ resistance to deformation and slows their shape-recovery.

**Figure 2:**
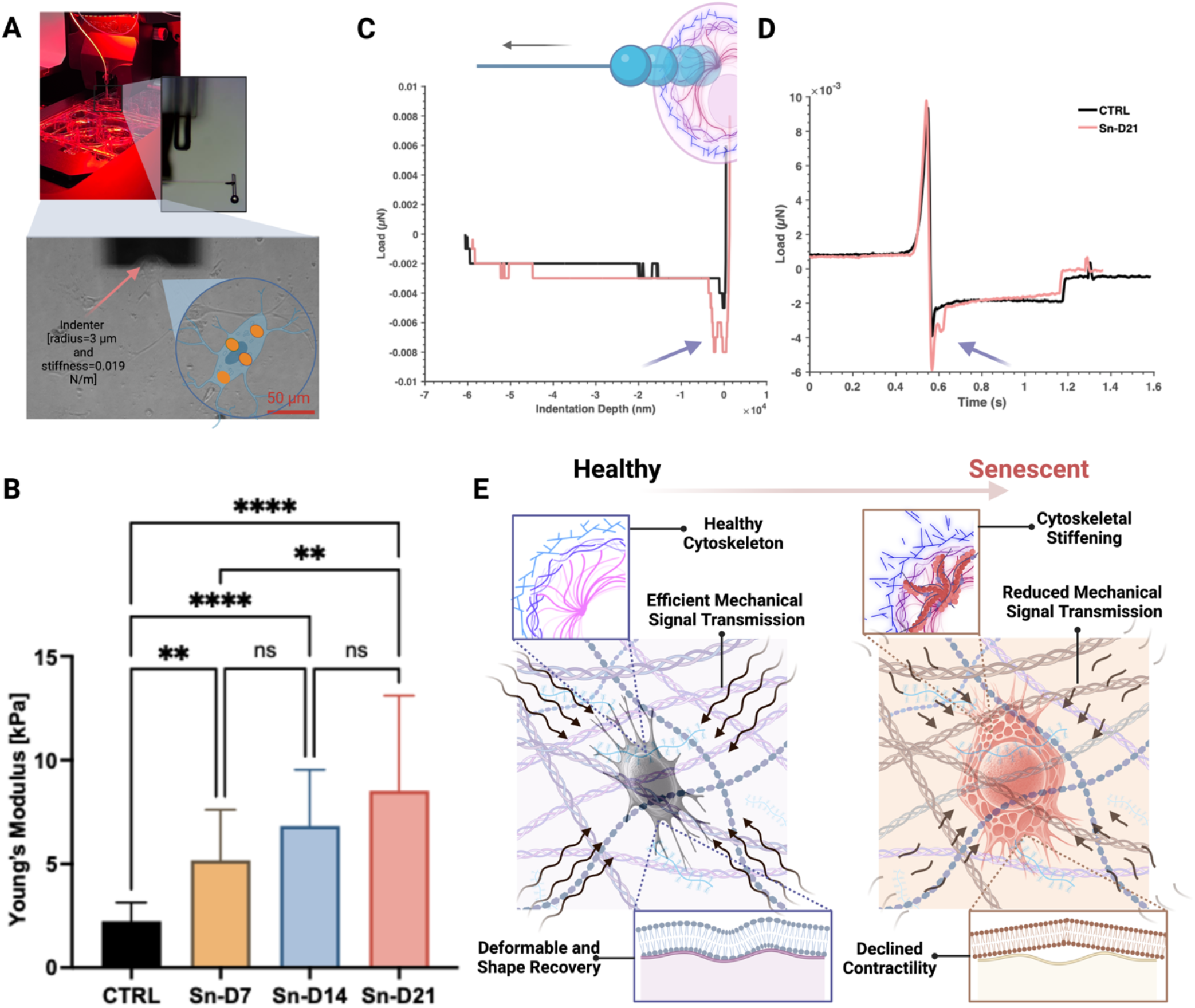
Mechanobiological Changes in Senescent Osteocytes. (A) Representative images of our mechanobiology experimental setup (B) Mechanical measurements showing increased Young’s modulus and potentially altered viscoelastic properties in senescent osteocytes. A minimum of N=90 single cells per study group (i.e., CTRL, Sn-D7, Sn-D14, and Sn-D21) were mechanically tested at every time point. Young’s modulus was obtained using Hertzian contact model with Poisson’s ratio ν = 0.5. (C) Mean unloading portions of indentation curves and (D) load-time curves, demonstrating significantly higher minimum adhesion force and delayed recovery in senescent cells (p=0.0084). In plot (C), all indentation data from control and Sn-D21 were used, while plot (D) includes experimental data from cells exhibiting osteocyte-like morphology. (E) Schematic drawing connecting these findings to the impaired mechanosensation and mechanotransduction capabilities of senescent osteocytes. These changes contribute to age-related bone loss and fragility. Data are presented as means ± SD, with statistical significance denoted by *p < 0.05, **p < 0.01, ***p < 0.001, and ****p < 0.0001.

These new experimental outcomes provide a foundational understanding of the mechanisms by which the mechano-responsiveness of osteocytes declines as they become senescent, potentially compromising their ability to sense and respond to mechanical stimuli (see Figure 2E). Probing these biophysical changes alongside gene expression variations (see Figure 1C) helped us to decipher biophysical implications of biomolecular changes in cellular senescence. Sost, an inhibitor of the Wnt/β-catenin signaling pathway, was notably upregulated at day 21 (Sn-D21). This may suggest that Sost might play a crucial role in cytoskeletal reorganization and the resultant increased stiffness through an indirect interaction with the Wnt/β-catenin pathway.^27,28^ It has been suggested that a divergent canonical Wnt pathway regulates the microtubule cytoskeleton to promote cell movement, indicating that Wnt signaling influences the cytoskeletal dynamics.^29^ According to a previous study,^30^ Wnt3a directly stimulates Mepe transcription via β-catenin and Lef-1 and indirectly through an autocrine BMP-2 loop. These pathways play crucial roles in bone homeostasis and remodeling. ^16,19,30^ Here, slight increase in expression level of Mepe observed on days 14 and 21 post-irradiation (Figure 1D-E) likely reflects adaptive response of primary osteocytes to irradiation, potentially contributing to cytoskeletal stiffening. Additionally, the decreasing expression trend of Mmp-9 from CTRL to Sn-D21 (see Figure 1D) could be linked to reduced ECM degradation. This reduction may contribute to a more rigid cytoskeletal structure in senescent cells.^31,32^ Thus, while Mepe and M-mp9 are two of the primary markers associated with mineralization and ECM remodeling, their altered expression in senescent cells underscores broader implications for cytoskeletal organization and mechanical properties.

The significant increase in p16^Ink4a^ expression level, a potent cyclin-dependent kinase inhibitor, has been previously linked to bone loss and age-related pathologies.^19,20,22,33,34^ Elevated p16^Ink4a^ levels are associated with the accumulation of senescent cells, which contribute to impaired tissue function and increased fracture risk in osteoporosis.^19,20,22,33,34^ The causal role of p16^high^ cells has been implicated in various pathological conditions, including osteoporosis.^4,21,22,35^ In Figure 1C, the observed decrease in p16^Ink4a^ expression at day 21 could reflect changes in the cell population or an adaptive response to irradiation. This variation may be due to the heterogeneous nature of the cell population in culture, leading to differential biophysical responses. Overall, these observations align with previous studies^19,20,22,33,34^ that have shown senescent osteocytes exhibit increased p16^Ink4a^ expression with age, contributing to reduced bone remodeling capacity.

To date, there are only two published reports that have investigated changes in the mechanical properties of senescent cells at the single-cell level, both focusing on fibroblasts and skin as their model systems.^36,37^ Therefore, the present study is pioneering in exploring senescence-induced changes in the mechanical properties of primary osteocytes, which are the most abundant mechanosensory cells in the human body. Previous studies^36,37^ have used either traction force microscopy (TFM) or atomic force microscopy (AFM). While these methods have contributed significantly to our understanding, they have limitations that our mechanobiology experimental setup overcomes. TFM’s reliance on the mechanics of the PDMS substrate for cellular force inference introduces complexities, while AFM, though direct in force application, cannot achieve the high specificity in indenting as well as our method. The use of a sharp diamond-shaped indenter in AFM can also affect outcomes in single-cell studies. Moreover, non-invasive and real-time monitoring techniques such as those used in this study can be adapted for real-time assessment of cell mechanics in living tissues. These methods allow biophysical markers to be assessed in live cells without the need for fixation or labeling, preserving the natural state of the cells and enabling longitudinal studies of senescence and aging. Overall, our approach enhances the ability to monitor biophysical markers of single cells over time, providing valuable insights into the dynamics of cellular aging.

In summary, the studied mechanobiological changes not only impair the mechanosensation and mechanotransduction capabilities of osteocytes but also contribute to age-related bone loss and fragility. The correlation between gene expression changes and mechanical properties underscores the importance of using such biophysical markers in conjunction with highly variable genomic markers to provide more reliable indicators of cellular senescence. Additionally, our new findings further support the hypothesis that biomechanical stimulation or exercise of cells could be used as a therapeutic approach to counteract the effects of aging on osteocytes. Specifically, by targeting the mechanobiological pathways involved in cytoskeletal stiffening, it may be possible to rejuvenate aging cells and improve bone health in elderly populations. An intriguing question for future research is whether these biophysical fingerprints (i.e., mechanical markers) could provide a better indication of whether the senescence state of a cell is beneficial or detrimental. This understanding could significantly influence how we perceive and manage cellular senescence in osteocytes and potentially other cell types within the musculoskeletal system.

## AUTHOR CONTRIBUTIONS

Maryam Tilton, Megan Weivoda, and James Kirkland designed the study. Maryam Tilton performed the experiments, researched, and analyzed the data. Maryam Tilton, Megan Weivoda, Maria Astudillo Potes, Anne Gingery, Alan Liu, Tamara Tchkonia, Lichun Lu, and James Kirkland provided resources and interpreted the data. Maryam Tilton wrote the original manuscript, and all co-authors reviewed and revised the manuscript. All authors read and approved the final manuscript.

