## Supplemental Materials for "Stiffening Symphony of Aging: How Senescent Osteocytes Lose Their Elastic Rhythm"

**Supplementary Methods**

**Primary osteocyte isolation, culture, and senescence induction:** Following the previously established protocol,[1–3] primary osteocytes were isolated from vertebrae of C57BL/6 WT mice (N=10; female). A highly enriched population of osteocytes were collected from digestion cycles 7 to 9, centrifuged, and seeded on 0.15 mg/mL rat tail type I collagen-coated plates (Fisher Scientific).[4] The culture medium comprised α-MEM (Sigma), enriched with L-glutamine, nucleosides, 5% heat-inactivated fetal bovine serum, and 5% calf serum (both from Sigma). After two passages to ensure cell purity and viability, the cells were cultured in type I collagen-coated 6-well plates. A subset of these cultures was exposed to 10 Gy of cesium irradiation (CellRad, Precision X-Ray) to induce senescence *in vitro*.[1,2] Each experimental condition was replicated across three wells.

**Senescence associated β-Galactosidase (SA-β-Gal) assay:** Following the previously established protocol,[5] cellular senescence was assessed by measuring SA-β-Gal activity, a reliable marker for senescent cells, using the CellSignaling SA-β-Gal staining kit, as per the manufacturer's instructions. The SA-β-Gal assay is currently the most widely used method for detecting senescence at the single-cell level due to its convenience and effectiveness. For quantitative analysis, ten randomly selected fields of view (both central and peripheral) were captured for each sample from each experimental group using brightfield inverted microscopy in a blinded manner. To quantify SA-β-Gal activity (%), an intensity-based analysis was conducted using ImageJ software. Brightfield microscopy images were converted to 8-bit grayscale to focus on the intensity of staining. A consistent threshold was applied to distinguish positive SA-β-Gal staining from the background, determined by the histogram of pixel intensity. The 'Analyze Particles' feature in ImageJ quantified the area stained relative to the total field area, providing a measure of SA-β-Gal activity as a percentage of the senescent cells.

**Real-Time quantitative polymerase chain reaction (RT-qPCR):** For RT-qPCR, biomarker tracking was systematically performed at various culture time-points (days 7, 14, and 21), utilizing a methodology extensively described in previous literature.[1–4] RNA isolation was achieved using Trizol reagent coupled with mechanical homogenization (i.e., ceramic bead mill tubes), followed by purification with the GeneJet RNA Purification Kit (Thermo Fisher). The RNA concentration was determined using the NanoDrop One spectrophotometer (Thermo Fisher). Subsequently, the purified RNA from each study group and time point was reverse transcribed to cDNA using the SuperScript IV VILO Master Mix (Thermo Fisher). In this pilot study, we focused on evaluating the expression levels of key biomarkers relevant to primary osteocyte function and senescence. Specifically, the expression of osteocyte-specific markers such as Matrix Extracellular Phosphoglycoprotein (MEPE) and Sclerostin (SOST), along with well-validated chronic senescence effectors in bone tissue like p16$^{Ink4a}$ and p21, and Matrix Metalloproteinases such as MMP9, was quantified. These targets were selected based on their established relevance to cellular senescence within the context of bone loss as validated in prior studies conducted by our team.[6–9] Detailed information regarding the Primer's used in these experiments is provided in Table 1. For quantifying mRNA levels, the $2^{-\Delta\Delta Ct}$ method was employed to calculate the normalized RNA expression for each target gene relative to the control gene.

*Table 1: List of Primers Used for RT-qPCR.*

| Genomic Target | Forward Primer Sequence | Reverse Primer Sequence |
|---|---|---|
| Sost | TCCTCCTGAGAACAACCA | CTGTACTCGGACACATCTT |
| Mepe | TGCTGCCCTCCTCAGAAATATC | GTTCGGCCCCAGTCACTAGA |
| *Cdkn2a (p16$^{Ink4a}$)* | GAACTCTTTCGGTCGTACCC | AGTTCGAATCTGCACCGTAGT |
| *Cdkn1a (p21)* | GAACATCTCAGGGCCGAAAA | TGCGCTTGGAGTGATAGAAATC |
| *Mmp9* | TGAGTCCGGCAGACAATCCT | CCCTGGATCTCAGCAATAGCA |
| Gapdh | ACTCAAGATTGTCAGCAAT | CCATCCACAGTCTTCTGGGT |
| L32 | CCATCTGTTTTACGGCATCATG | TGAACTTCTTGGTCCTCTTTTTGA |

**Immunofluorescence (IF) Staining:** To visualize the cytoskeleton during single-cell mechanical testing (i.e., nanoindentation), CellMask Deep Red Actin Tracking Stain (Thermo Fisher) was employed on live cells. Post-nanoindentation, cells underwent IF staining for F-actin and nuclei. The protocol involved washing the cells thrice with DPBS, fixing with 4% paraformaldehyde (PFA), and subsequent staining using CellMask for actin and DAPI for nuclei. Samples were imaged using our inverted fluorescence microscope on the Pavone system with 20x (NA=0.5) objective.

**Single cell mechanical testing:** Mechanical properties of the primary cells (2D culture) were measured at sub-cellular resolution using optical fiber-based interferometry nanoindenter (Pavone; Optics11Life). This instrument enabled single-cells mechanical characterization in live culture conditions. For these experiments, we used a spherical probe with R=3 μm and 0.019 N/m stiffness. The peak load threshold was set at 0.01 μN. A minimum of 30 indentation curves were obtained from each culture well at different locations (i.e., central and peripheral). We used Hertzian contact mechanics model[10–12] with our load-indentation data to determine the Young's modulus.

**Statistical analyses:** All statistical analyses were conducted using Prism9 software. We employed one-way and two-way ANOVA to analyze biophysical data (i.e., Young's Modulus) and biomarker expressions, respectively. Post-ANOVA, Tukey's multiple comparison post hoc test was performed to determine significant differences among experimental groups.
